# E(3) Invariant Representations of Biomolecules

**DOI:** 10.1101/2025.03.26.645520

**Authors:** Julio Candanedo

## Abstract

We present a coordinate-independent method for encoding protein structures using anchor-point-based trilateration that respects Euclidean symmetry while avoiding 𝒪(*N*^2^) scaling problems of distance matrices. Our approach achieves near-perfect reconstruction of protein structures, and when combined with dimensionality reduction techniques, effectively captures conformational dynamics in molecular trajectories.

## 1 Introduction

Protein structure determination is fundamental to understanding biological function, guiding drug discovery, and elucidating disease mechanisms. While experimental techniques such as X-ray crystallography, NMR spectroscopy, and cryo-electron microscopy have revolutionized our ability to resolve biomolecular structures, the computational representation and analysis of these structures present unique challenges. Proteins and other biomolecules are typically represented as collections of atoms in three-dimensional space, with coordinates stored in a matrix format. However, this representation introduces a fundamental complication: the same physical structure can be represented by infinitely many different coordinate sets due to arbitrary choices of reference frame. This coordinate frame dependence creates significant obstacles for structural comparison, trajectory analysis, and machine learning applications that rely on consistent data representations. In this paper, we address this challenge by developing a coordinate-independent encoding scheme that respects the inherent symmetries of three-dimensional space while maintaining computational efficiency.

The structure of protein and biomolecules, may be represented by an atomistic model consisting of atom types and their corresponding coordinates. These coordinates may be compiled into a 2-index matrix *R*_*ix*_ ∈ *ℝ*^*N ×d*^, with index *i* enumerating *N* atoms and index *x* enumerating each atom’s Cartesian coordinates (typically *d* = 3). Unfortunately, this representation is not unique; e.g., orthogonal-group rotations given by a matrix product 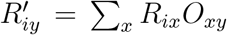, give rise to coordinates with new numerical values. In addition to rotations, we can have translations 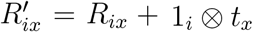, for some translation given by the 3-dimensional vector *t*_*x*_. Together, these transformations, *isometries*, define the Euclidean Group, symbolically represented as *E*(3). Moreover, all of these infinitely possible coordinates are valid representations of our biomolecular structure. Therefore, during the analysis of the biomolecular structural data, it is important to know whether two data entries for the biomolecular structure actually include the *same* structure (with different coordinates). Using the raw molecular coordinates therefore is unlikely to be suitable for molecular dynamics and structural biology analysis schemes.

Our first task is therefore to determine whether two sets of coordinates *R*_*ix*_ and 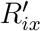 define the same structure. This is a routine algorithm, known as the Kabsch-Umeyama algorithm Kabsch (1976); Umeyama (1976), and what we wish to discover is the rotation-matrix and translation vector *{O*_*xy*_, *t*_*y*_*}* as before (with ⊗denoting the tensor/outer product):

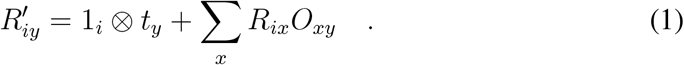

A subtlety is that the Kabsch-Umeyama algorithm does not take into account reflections, i.e., the rotation matrix *O*_*xy*_ is an element of the special orthogonal group (not the full orthogonal group). This is easily addressed by the sign of the determinant det (*O*_*xy*_).

A known structural invariant of E(3) is the entire distance matrix *D*_*ij*_ between all atoms. That is, for an arbitrary E3 transformation of our coordinates, the distance matrix is exactly the same:

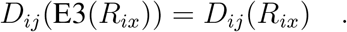

This implies that every matrix element of our distance matrix is an E3 invariant scalar. Therefore, the entire distance matrix can be used for data analysis, implicitly knowing if two structures are the same. A major issue with this approach is the scaling 𝒪(*N*^2^) in both computation and memory. Instead of this, as we mentioned, a subset of matrix elements is also E(3)-invariant. In fact, any transformation of the distance-matrix is also E(3) invariant.

If we have the complete distance matrix, we can obtain the coordinates in their entirety via the spectrum of the Gram Matrix, defined by (with δ_*ij*_ the Kronecker-delta/identity matrix, and 1_*ij*_ an *N × N* matrix with all entries equal to 1, and ^⊙2^ denoting the element-wise power):

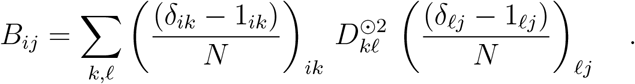

Yielding the following coordinates:

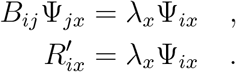

The Euclidean square distances may be computed via the law-of-cosines:

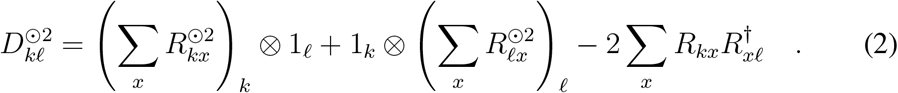

Although the Gram matrix is of size *N × N*, its true rank is at most the dimensionality *d* = 3. Therefore, most of the matrix elements in our distance and Gram matrix encode redundant information. Considering developments in matrix completion, it was surmised that a stochastic sparse subsampling of order 𝒪(3*N*^6/5^ log *N*) might be sufficient to encode this information Candes and Recht (2009). The idea is to stochastically sample *B*_*ij*_ and cast this sampling into a sparse matrix, and then use matrix factorization to obtain the coordinates *U*_*ix*_, i.e., a fitting procedure using the following loss function: 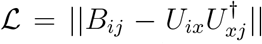. However, the matrix type is not suitable for most matrix factorization schemes, as the distribution of matrix elements spans many orders of magnitude and has a skewed Maxwell-Boltzmann distribution.

However, there exists a simpler solution! It was noted that 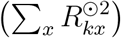 in Eq. 2 actually encodes the distances from a special point, the origin. If we fixed the 0th point (xor the 1st depending on who is counting) to the origin, 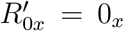, this quantity would actually encode the 0th row of the distance matrix: *D*_0*i*_. But why stop there? What if we selected *A* number of so-called *anchor-points*; therefore we use *trilateration* to reconstruct the entire distance matrix from a subset of the entire 𝒪(*N*^2^) elements. For clarity, we expand our notation: new indices *a, b* enumerate the anchor points with maximum size *A*, and *p, q* enumerate only non-anchor points with maximum value of *M*. Together, anchor and non-anchor points define our system *N* = *A* + *M*, enumerated by indices *i, j*. Therefore, we define an E(3) invariant encoding as part of the distance matrix:

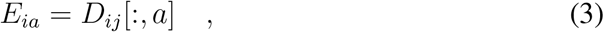

with the matrix slicing ([:, *a*]) in the numpy convention, obtaining only *a* elements of the entire row. Now we show that *E*_*ia*_ is able to fully reconstruct the entire distance matrix.

Next, using *E*_*ia*_, we would like to return to the concrete coordinates. The anchor points have pairwise distances fully explicitly computed. Therefore, we can extract a distance matrix:

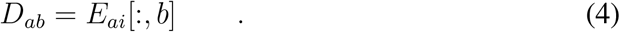

This matrix, *D*_*ab*_, may be converted into a Gram matrix and subsequently diagonalized to obtain the coordinates of the anchor points; this is denoted by *W*_*ax*_. Using this result and a special anchor-point, an anchor anchor-point, without loss of generality let’s suppose this is the point with index ‘*a* = 0’. Then we can define the following intra-anchor matrix:

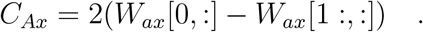

with the new index *A* being index *a* excluding that special point. We next construct the pseudo-inverse of *C*_*Ax*_; this can be achieved with Singular-Value-Decomposition (SVD), inverting the singular values, and subsequently reconstructing the resulting matrix, yielding 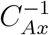.

Next, we construct a matrix:

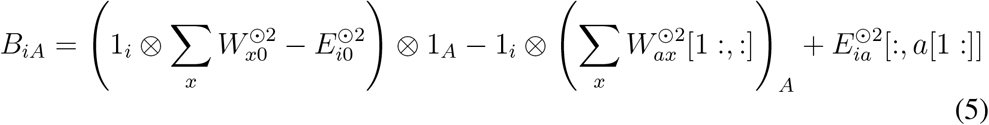

Finally, our reconstructed coordinate come:

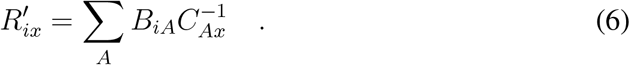

With 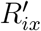 in hand, the entire distance-matrix may be reconstructed, if desired, using only the information encoded in *E*_*ia*_!

## 2 Application to Various Proteins

Let us now consider this algorithm in action to encode and decode real-life experimental results for protein structure. For example, let us consider the following Protein-Data-Base (PDB) entries: `1DLL` by Emsley et al. (2000), `1DYL` by Mancini et al. (2000), `6PFY` by Gisriel et al. (2019), and `1C2W` by Mueller et al. (2000). Once downloaded from the PDB, using `ProDy` *et al*. (2011), we encode and subsequently decode the original coordinates *R*_*ix*_ to obtain a new set of coordinates *P*_*ix*_. We then use a custom Kabsch-Umeyama implementation to align the decoded coordinates *P*_*ix*_ with the original coordinates *R*_*ix*_ (taking into account reflections). The results of our reconstruction comparisons are shown in Fig. 1. All these plots (rendered by 3Dmol et al. (2015)) show two cartoon structures, one in purple (slightly transparent) and the other in yellow. All reconstructed structures have an RMSD of 10^−11^ Å and are so close to the original that the two structures in Fig. 1 appear as one with a mixed mustard color.

**Figure 1:**
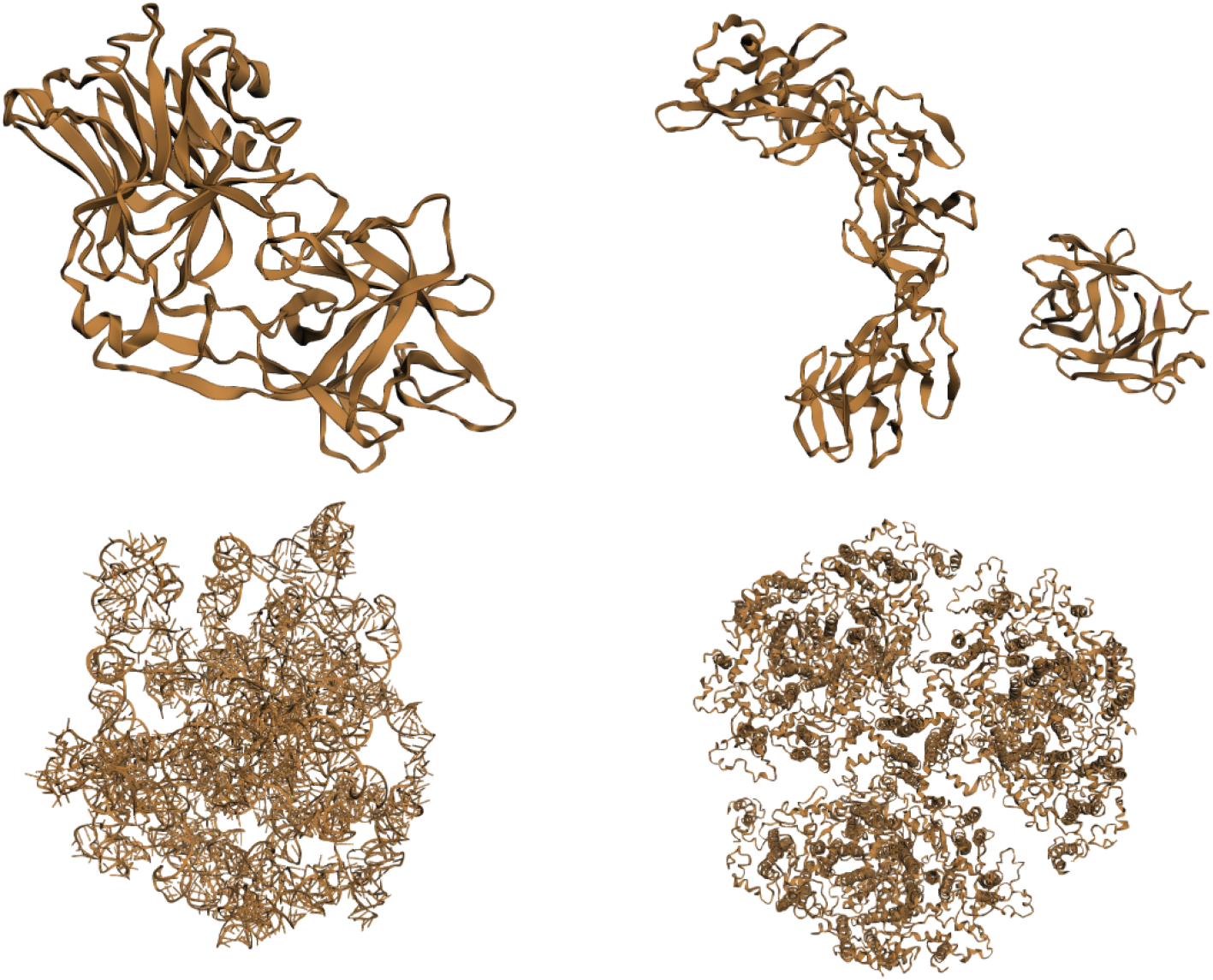
Above from upper-left to lower-right (in a “z” shape) are PDB structures: 1DLL, 1DYL, 1C2W, and 6PFY. On each plot are two structures: one purple and one yellow with opacity on the purple structure.

## 3 Application to a Trajectory

In this section, we consider a series of protein configurations, which we call a trajectory *R*_*tix*_, a 3-dimensional tensor with the time index *t* for *T* time-steps. For each frame, at each *t*, we may obtain an E3-encoded structure, that is, *E*_*tia*_. This dataset may be subsequently reshaped into a matrix *E*_*tX*_. The dimension of *X* is very large 𝒪(*NA*), and therefore we opt to reduce the dimensionality of the structure using the diffusion map algorithm Coifman et al. (2005). This would result in a compressed trajectory *E*_*tα*_. To demonstrate this, we used a Molecular dynamics (MD) trajectory dataset produced by Fan (2019); Hussein et al. (2023) on the YiiP membrane protein^2^, and processed it using a custom diffusion map code. We use the following pipeline, starting from the raw downloaded trajectory tensor 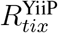:

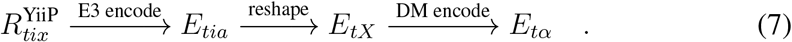

In addition to keeping track of the structure, we can also keep track of the order of the trajectory using colors (red for structures closest to time 0, violet for most distant times, and a rainbow spectrum in between). Our results are plotted in Fig. 2, and the diffusion map allows instant visualization of a clear path that follows from the initial state (red) to the final state (violet), with the order of structures in the original trajectory intact.

**Figure 2:**
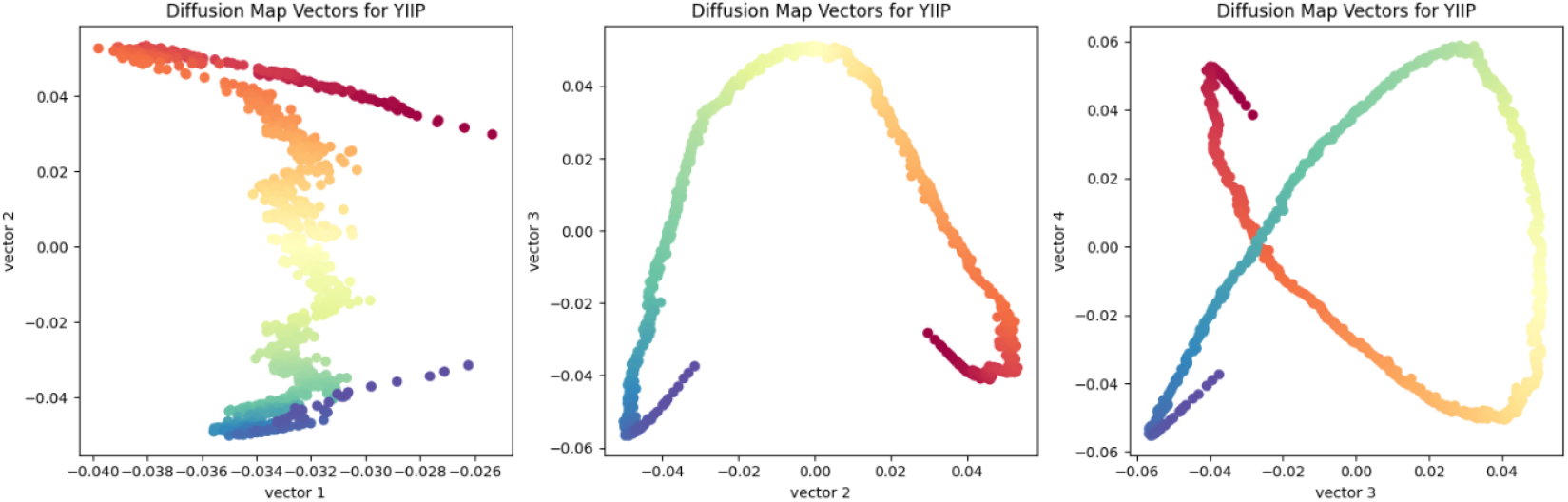
Above are 3 plots with the diffusion map embeddings (or vectors) plotted against each other to show the parametric structure of our MD trajectory. Each point is colored according to its original trajectory order (the initial time is colored in red, later times are in violet).

## 4 Concluding Remarks

The algorithm demonstrated here provides an efficient and mathematically rigorous method for encoding biomolecular structures while properly respecting the Euclidean group symmetries. By using anchor-point based trilateration, we achieve several key advantages: Near perfect structural reconstruction without the 𝒪(*N*^2^) memory and computational scaling issues associated with full distance matrices.

Coordinate-independent representation that facilitates direct comparison between structures. Compatibility with advanced dimensionality reduction techniques, as demonstrated with diffusion map analysis. These properties make our approach particularly valuable for analyzing large structural datasets, including molecular dynamics trajectories where *E*(3) invariance is essential. Our brief demonstration with the YiiP membrane protein trajectory illustrates how this encoding can reveal conformational pathways while preserving the temporal ordering of states. Future work could extend this approach in several directions. First, incorporating atom type information alongside geometric encoding could enhance chemical specificity. Second, this representation could be used for machine learning applications, particularly in generative models for biomolecular design where the *E*(3) invariance is critical. Third, the anchor-point selection strategy could be optimized based on structural properties rather than random sampling, potentially improving encoding efficiency for specific biomolecular classes. Finally, this approach could be extended to analyze structural ensembles from cryo-EM or NMR experiments where multiple conformations are simultaneously present.

## A. JAX implementation of algorithm

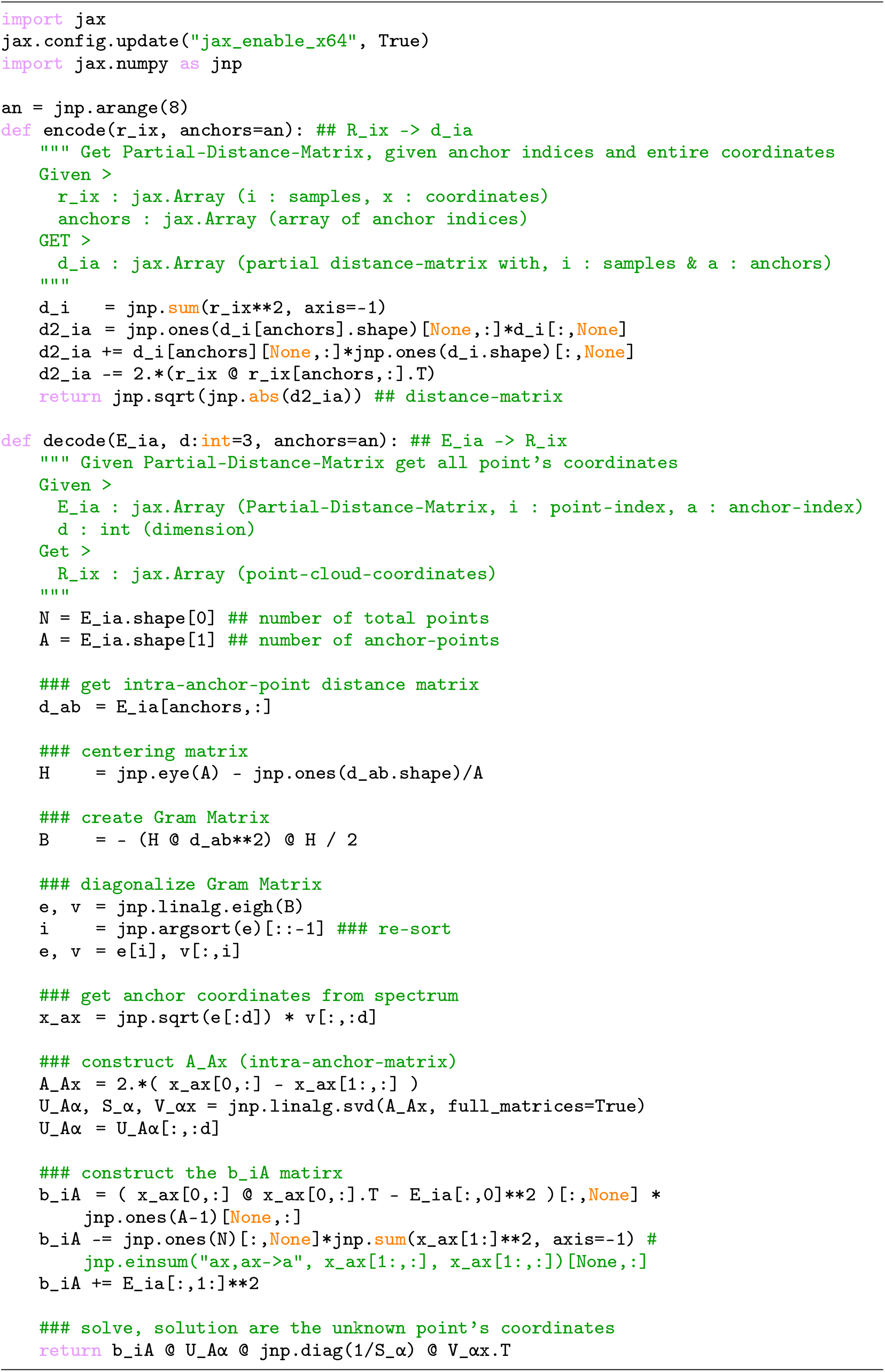

## Footnotes

2 ”Molecular dynamics (MD) trajectory files of the YiiP membrane protein in a POPE:POPG 4:1 model membrane. The equilibrium simulation was performed in the NPT ensemble at T=300K and P=1 bar. The system was simulated with Gromacs 2018.1, using the CHARMM36 force field, the TIP3P explicit water model, NaCl at approximately 100 mM concentration, and Zinc ions. Trajectory frames were saved every 100 ps for a total of 9 ns and 90ns simulated time. The topology contains the whole system (protein, membrane, water and ions).” Fan (2019).

## Notes

### Competing Interest Statement

The authors have declared no competing interest.

## References

Candes, E. and Recht, B. (2009). FoCM, 9:717–772. 0805.4471.

Coifman, R. R., Lafon, S., Lee, A. B., Maggioni, M., Nadler, B., Warner, F., and Zucker, S. W. (2005). PNAS, 102(21):7426–7431.

Emsley, P., Fotinou, C., Black, I., Fairweather, N. F., Charles, I. G., Watts, C., Hewitt, E., and Isaacs, N. W. (2000). JBC, 275:8889–8894.

Fan Shujie Beckstein, O. (2019). figshare dataset.

Gisriel, C., Coe, J., Letrun, R., Yefanov, O. M., Luna-Chavez, C., Stander, N. E., Lisova, S., Mariani, V., Kuhn, M., Aplin, S., Grant, T. D., Dörner, K., Sato, T., Echelmeier, A., and Jorvani Cruz Villarreal, e. a. (2019). Nature Communications.

Hussein, A., Fan, S., Lopez-Redondo, M., Kenney, I., Zhang, X., Beckstein, O., and Stokes, D. L. (2023). eLife, 12:RP87167.

Kabsch, W. (1976). Acta Cryst.

Mancini, E. J., Clarke, M., Gowen, B. E., Rutten, T., and Fuller, S. D. (2000). Molecular Cell, 5:255–266.

Mueller, F., Sommer, I., Baranov, P., Matadeen, R., Stoldt, M., Wöhnert, J., Görlach, M., van Heel, M., and Brimacombe, R. (2000). Journal of Molecular Biology, 298(1):35–59.

Dmol Rego, N., Koes, D., and Notes, A. (2015). Bioinformatics, 31:1322–1324.

ProDy, AB., LM, M., and I., B. (2011). Bioinformatics, 27:1575–1577.

Umeyama, S. (1976). IEEE PAMI.

